# A Mathematical Model for the Immune-Mediated Theory of Metastasis

**DOI:** 10.1101/565531

**Authors:** Adam Rhodes, Thomas Hillen

## Abstract

Accumulating experimental and clinical evidence suggest that the immune response to cancer is not exclusively *anti*-tumor. Indeed, the *pro*-tumor roles of the immune system — as suppliers of growth and pro-angiogenic factors or defenses against cytotoxic immune attacks, for example — have been long appreciated, but relatively few theoretical works have considered their effects. Inspired by the recently proposed “immune-mediated” theory of metastasis, we develop a mathematical model for tumor-immune interactions at two anatomically distant sites, which includes both *anti*-and *pro*-tumor immune effects, and the experimentally observed tumor-induced phenotypic plasticity of immune cells (tumor “education” of the immune cells). Upon confrontation of our model to experimental data, we use it to evaluate the implications of the immune-mediated theory of metastasis. We find that tumor education of immune cells may explain the relatively poor performance of immunotherapies, and that many metastatic phenomena, including metastatic blow-up, dormancy, and metastasis to sites of injury, can be explained by the immune-mediated theory of metastasis. Our results suggest that further work is warranted to fully elucidate the protumor effects of the immune system in metastatic cancer.

## 1. Introduction

Although metastasis is implicated in over 90% of all cancer related deaths (Gupta and Massague, 2006; Liu and Cao, 2016; Valastyan and Weinberg, 2011), a full understanding of the process remains elusive. Accumulating evidence, including the observation that patients who received peritoneovenous shunts that inadvertently released large numbers of cancer cells directly into the patients’ blood stream saw no increased rate of metastasis (Tarin et al., 1984), and the immune-mediated preparation of the premetastatic niche (PMN) by Kaplan and collaborators (Kaplan et al., 2005), has brought into question the prevailing view of metastasis as a passive, random process. Of particular interest is the recent hypothesis that the immune system — in addition to its well-known *anti*-tumor role — plays an active *pro*-tumor role in metastatic disease Cohen et al. (2015); de Mingo Pulido and Ruffell (2016); Shahriyari (2016). Well supported by experimental and clinical observations, this hypothesis and its consequences has yet to be fully investigated. The goal of the present work is to begin this investigation through the development and analysis of a mathematical model for the immune-mediated theory of metastatic cancer. In the following section, we briefly highlight the biological evidence for this theory to justify our mathematical model, which is introduced in Section 2. We also include a short discussion of previous mathematical models of metastasis in order to contrast them against our approach.

### 1.1. The Immune-Mediated Theory of Metastasis

A link between the immune system and cancer has been noted for a long time (Balkwill and Coussens, 2004; Walter et al., 2011), with investigators referring to tumors as “wounds that do not heal” (Dvorak, 1986, 2015) or suggesting that they are the result of an uncontrolled healing process (Meng et al., 2012). Recently, “avoiding immune destruction” and “tumor-promoting inflammation” were identified as an emerging hallmark and an enabling characteristic of cancer, respectively (Hanahan and Weinberg, 2011). More specifically, a number of authors have synthesized the accumulating evidence implicating the immune system in metastasis to formulate the immune-mediated theory of metastasis (Cohen et al., 2015; Shahriyari, 2016). In this section we present a brief summary of the relevant evidence to support this theory organized using the “metastatic cascade” framework. Within the well-used metastatic cascade framework, metastasis is seen as a sequence of biological processes beginning with the development, growth, and local invasion of a primary tumor, and followed by the preparation of a premetastatic niche, entrance into, travel through, and exit from the vascular system, and concluding with the growth and development of a secondary, metastatic tumor. The metastatic cascade is depicted in Figure 1, with special attention paid to the immune effects at each step. We now highlight the specific immune cells involved at each step of the metastatic cascade and outline their roles.

**Figure 1:**
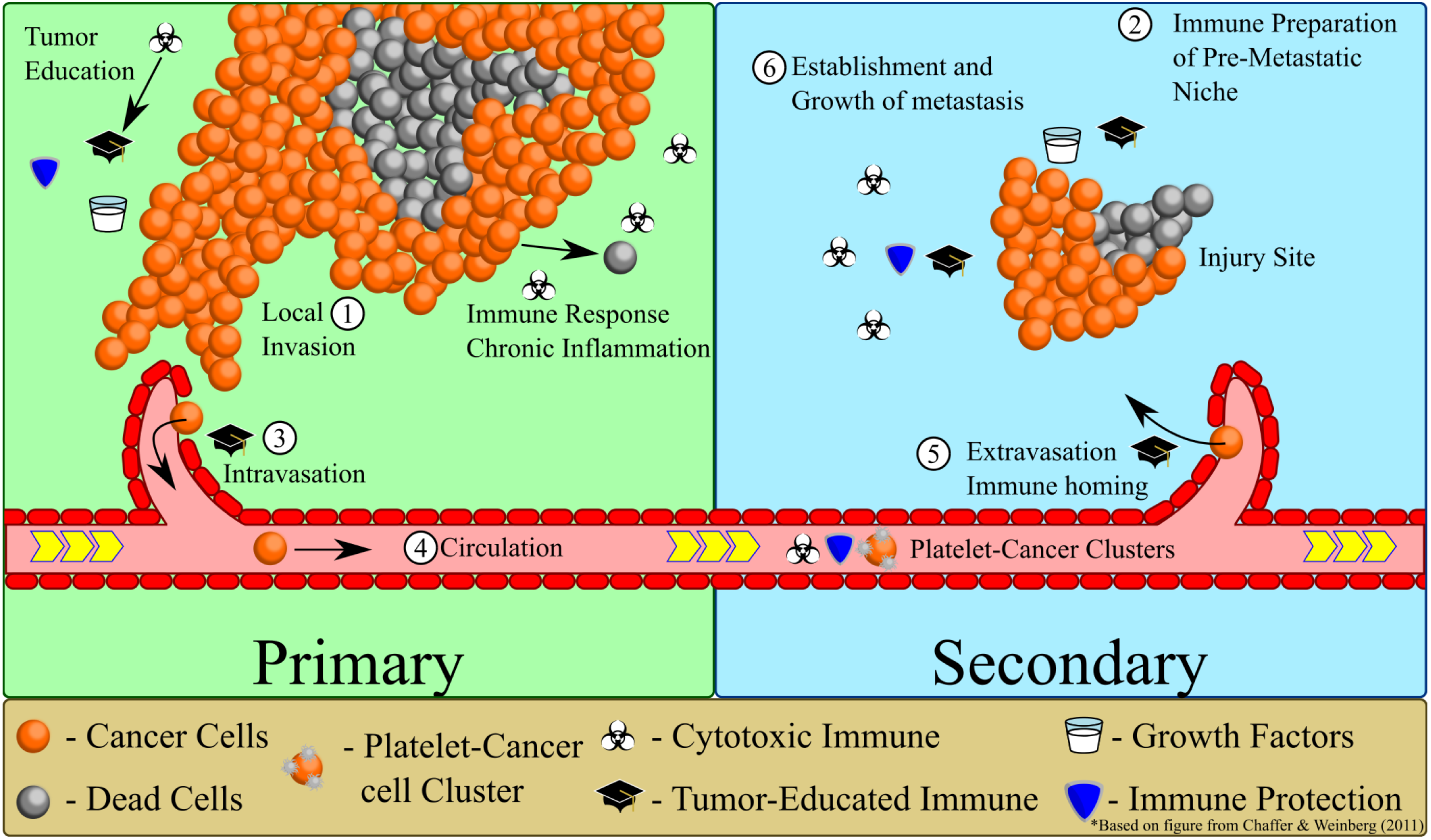
Cartoon model of the immune-mediated model of metastasis. Based on figure from Chaffer and Weinberg (2011).

#### Step (1): Primary Tumor Growth and Local Invasion

Before cancer can spread throughout the body, an initial *primary* tumor must first develop (see Figure 1, (1)). Immune involvement in this stage of metastasis has long been acknowledged (Dvorak, 1986; Hanahan and Weinberg, 2011). A large amount of research suggests that the role of the immune system in tumor progression is hardly straightforward (de Mingo Pulido and Ruffell, 2016; Erdman and Poutahidis, 2010). Indeed, while the *anti*-tumor roles of the immune system are well known — the cytotoxic effects of natural killer (NK) cells, “classically activated” M1 macrophages, and CD8^+^ T cells, for example (Joyce and Pollard, 2009) — many immune cells have also been shown to play *pro*-tumor roles in primary tumor development (de Mingo Pulido and Ruffell, 2016; Joyce and Pollard, 2009). For example, regulatory T cells (T_regs_), T helper type 2 (Th_2_) cells, neutrophils, and “alternatively” activated (M2) macrophages can all promote growth through inhibition of cytotoxic immune responses (Joyce and Pollard, 2009) or promotion of angiogenesis (de Mingo Pulido and Ruffell, 2016). For extensive reviews of the specific roles played by different immune cells, please see the reviews by de Mingo Pulido and Ruffell (2016) and Joyce and Pollard (2009).

In addition to the contradictory anti-and pro-tumor roles played by immune cells, there is evidence suggesting that tumors can “convert” or “educate” *anti*-tumor cytotoxic (CT) immune cells into *pro*-tumor immune cells (Oleinika et al., 2013). Shahriyari (2016) has proposed that, at sites of chronic inflammation, the local immune cells become adapted to the wound healing process, resulting in increased proliferative signaling and decreased cytotoxic activity. Liu et al. (2007) have demonstrated that tumor-derived transforming growth factor (TGF) *β*, derived from the murine prostate tumor TRAMP-C2 and renal cell carcinoma RENCA, can induce the transition of *anti* –tumor CD4^+^CD25^-^ T cells into *pro*-tumor CD4^+^CD25^+^ T_regs_. Such results allow for the notion of “tumor educated” (TE) immune cells (Liu and Cao, 2016); a term that will be used throughout this paper. This experimentally-validated notion of phenotypic plasticity amongst sub populations of immune cells is not entirely new, and has been considered for some time in the context of macrophages, with a continuum between *anti*-tumor M1 macrophages and *pro*-tumor M2 macrophages being proposed (Balkwill and Coussens, 2004; den Breems and Eftimie, 2016).

#### Step (2): Preparation of the Pre-Metastatic Niche

Often, metas-tatic dissemination is viewed as a passive process in which cancer cells shed from the primary tumor establish metastatic tumors at sites “downstream” of the primary tumor, in locations that the circulating tumor cells (CTCs) become stuck in small vessels (Chaffer et al., 2011; Hiratsuka et al., 2006). It has been shown, however, that this model of metastasis can only account for approximately 66% of all observed patterns of metastasis (Chambers et al., 2002), suggesting that there are additional factors to consider. In an update of Paget’s classic “seed and soil” hypothesis (Paget, 1989), the concept of a pre-metastatic niche (PMN) has been developed by several investigators. While the precise definition of a PMN is still being debated (Qian and Pollard, 2010), the key concept is that the PMN is a supportive *setting* in which metastatic tumors can more efficiently establish themselves, and which may (Dos Anjos Pultz et al., 2017) or may not be influenced by the primary tumor itself.

Of particular interest is the implication of the immune system in the development of the PMN. Numerous cells, proteins, and factors have been implicated in the preparation of the PMN, ranging from (primary tumor associated) vascular endothelial growth factor (VEGF)-A, tumor necrosis factor (TNF) *α* and TGF *β* (Liu and Cao, 2016), to immuno-attractant S100 proteins (Joyce and Pollard, 2009; Kitamura et al., 2015; Qian and Pollard, 2010) and matrix-degrading MMPs (Kitamura et al., 2015), to bone marrow derived cells (BMDCs) (Coughlin and Murray, 2010; Joyce and Pollard, 2009; Kaplan et al., 2005) and platelets (Joyce and Pollard, 2009; Shahriyari, 2016) (which can produce their own *pro*-tumor factors). The work of Kaplan and collaborators (Kaplan et al., 2005) showed that, not only did BMDCs arrive at the site of future metastasis *prior* to the arrival of any cancer cells, but once cancer cells *did* arrive, they localized to regions of high BMDC density, suggesting a supportive role for immune cells in metastatic establishment. Further implication of the immune system in metastatic establishment comes from Shahriyari (2016), who has suggested that wound healing sites, which are naturally populated with immune cells producing growth promoting and CT immune inhibiting factors, may act as a metastasis-supporting PMN, thereby providing a possible explanation for observations of metastasis to sites of injury (Kumar and Manjunatha, 2013).

Taken together, such results suggest a supportive role for the immune system in metastatic establishment, wherein the immune cells may aid in successful establishment of newly arrived cancer cell(s) by supplying growth factors (ex: platelets secreting pro-growth and angiogenic factors such as stromal-derived factor (SDF) 1 (de Mingo Pulido and Ruffell, 2016)) and protection from CT immune cells (ex: T_regs_ or adapted immune cells (Shahriyari, 2016)) (Figure 1 (2)).

#### Step (3): Intravasation

In order to establish a secondary tumor at an anatomically distant location from the primary tumor, cancer cells must travel from the primary site to the secondary site. Although cancer cells can be found in lymph nodes, it is believed that the major method of distant dissemination is through the vascular system rather than the lymphatic system (Chambers et al., 2002; Joyce and Pollard, 2009). In order to gain access to the vascular system, cancer cells, or small clusters of cancer cells (Friedl and Mayor, 2017), must leave the parenchyma and enter a blood vessel in a process called *intravasation*. While the precise mechanism underlying intravasation remains obscure, tumor-associated macrophages (TAMs) have been implicated in the process. In fact, specific studies have reported that intravasation occurred only where perivascular TAMs were located (see Joyce and Pollard (2009); Liu and Cao (2016) and references therein) (see Figure 1 (3)).

#### Step (4): Circulation

Upon entrance into the blood vessel, the cancer cells are subject to a litany of new dangers, including shear forces and immune defenses (see Figure 1 (4)). It is believed that platelets play a critical role in the protection of the cancer cell clusters while in circulation. Not only can they protect from the effects of shear force by forming clumps with the cancer cells, they may also act as shields against cytotoxic immune attack from NK cells (Joyce and Pollard, 2009; Kitamura et al., 2015). While the precise role of platelets is still debated (Coupland et al., 2012; Shahriyari, 2016), it has been shown that treatments with anti-coagulant and non-steroidal anti-inflammatory drugs (NSAIDs) can significantly decrease the rates of metastasis (Joyce and Pollard, 2009; Marx, 2004).

#### Step (5): Extravasation

It has been estimated that a primary tumor can shed tens of thousands of cells into the vasculature every day (Weiss, 1990). Experimental models of metastasis suggest that upwards of 80% of all those cells shed will successfully exit from the blood vessel (extravasate) at a distant secondary site (Cameron et al., 2000; Luzzi et al., 1998). As is the case with intravasation, macrophages have been implicated in the reverse process of extravasation (Kitamura et al., 2015; Liu and Cao, 2016; Qian and Pollard, 2010). In addition to the survival and growth factors (ex: TGF*β*, CCL2, VEGF-A) supplied by metastasis-associated macrophages (MAMs) as the tumor cells work to exit the vessel and enter the surrounding parenchyma (see Figure 1 (5)), tumor-MAM contact has also been shown to aid in cancer cell movement *through* the vessel wall. Platelets have also been shown to play a protumor role in this setting (Kitamura et al., 2015; Shahriyari, 2016), however they are not necessary for successful extravasation (Coupland et al., 2012). Another immune cell type that has been implicated in metastatic disease are neutrophils (Demers et al., 2012; Park et al., 2016) through the use of neutrophil extracellular traps (NETs) which can trap circulating tumor cells at a distant, hospitable site, or even increase local vascular permeability, allowing for easier extravasation of cancer cells into the surrounding parenchyma.

#### Step (6): Metastatic Establishment

Even though a large majority of cells shed from the primary tumor will successfully extravasate at a secondary site, less than 0.01% of them will successfully colonize a macroscopic metastatic tumor (Cameron et al., 2000; Luzzi et al., 1998) (see Figure 1 (6)). The experimental results of Cameron and colleagues (Cameron et al., 2000; Luzzi et al., 1998) suggest that this precipitous drop in survival occurs after the cells transition from quiescent to proliferative states, whereby they become more vulnerable to local defenses. While this transition can occur relatively soon after initial metastatic seeding of the secondary site, it is often the case that the newly arrived cancer cells lay dormant for an extended period of time before entering a proliferative phase (Hanahan and Weinberg, 2011). A possible explanation for the low efficiency of establishment observed may be found in an effective CT immune response (Eikenberry et al., 2009). However, the immune system plays contradictory roles in this step of metastasis, with a *pro*-tumor response mediated by BMDCs (Hanahan and Weinberg, 2011; Joyce and Pollard, 2009) or MAMs (Kitamura et al., 2015), which provide survival and proliferation signals, or inflammatory stromal cells (Joyce and Pollard, 2009), which provide protection from the cytotoxic effects of NK cells. Additionally, immune preparation of the PMN (see previous section) may also support metastatic development, and similar *pro*-tumor immune effects on growth and development may be common between primary and secondary sites.

### 1.2 Previous Mathematical Models of Metastasis

Metastasis, with its multi-step complexity and apparent stochasticity, is relatively difficult to study experimentally. Consequently, there is a great deal of uncertainty concerning the underlying dynamics of the process. Theoretical and mathematical models of the process are therefore of significant interest as they allow for detailed theoretical investigations of the underlying processes in order to test hypotheses and guide future biological research. In this section, we present a brief summary of the most relevant mathematical descriptions of metastasis that have been previously investigated.

Focusing on the supposed stochasticity of the process, many authors have developed stochastic models for cancer metastasis. From a stochastic modeling framework, Liotta et al. (1977) derived an expression for the probability of being *metastasis-free* as a function of time from primary tumor implantation. The Michor lab has spent significant effort investigating stochastic models for the emergence of the *metastatic phenotype* (Haeno and Michor, 2010; Michor et al., 2006). The *natural history of cancer* — that is, determining dates of disease initiation, first metastasis inception, etc. from clinical data — is the focus of the stochastic models emerging from the Hanin group (Hanin et al., 2006; Hanin and Rose, 2018). Recently, Frei et al. (2018) introduced a spatial model for cancer metastasis that takes the form of a branching stochastic process with settlement, providing one of the first models which explicitly accounts for travel between metastatic sites.

Based on their ease of analysis in comparison to stochastic models, many investigators have chosen to analyze deterministic models of metastasis. Saidel et al. (1976) introduced one of the earliest models of metastasis in the 1970s, using a simple compartmental ordinary differential equation (ODE) model that took into account the different steps in the metastatic cascade. More recently, Iwata (Iwata et al., 2000) introduced a partial differential equation (PDE) model describing the colony size distribution of metastases that takes the form of a transport equation subject to a non-local boundary condition. The Iwata model has since been adapted, analyzed, and confronted to data by several investigators — notably Benzekry and colleagues (Baratchart et al., 2015; Benzekry, 2011; Benzekry et al., 2014, 2017) — and has provided important insights into, among others, the effects of primary resection on metastatic tumor growth. To investigate the role of immune cell trafficking between metastatic sites and the so-called *abscopal effect* — in which cytotoxic treatment at one tumor site elicits an effect at a secondary site — the Enderling group has developed a model for tumor-immune interactions at multiple sites (Poleszczuk et al., 2016, 2017; Walker et al., 2018, 2017). While these works provide insight regarding tumor-immune dynamics in the metastatic setting, they are unable to provide details of the metastatic process itself. Franßen et al. (2018), on the other hand, have developed a multi-site model with spatially explicit dynamics at each of the sites that successfully captures the steps of the metastatic cascade.

Our model builds on Kuznetsov’s tumor-immune model (Kuznetsov et al., 1994) which has been the starting point for several investigators (Kuznetsov and Knott, 2001; Poleszczuk et al., 2016; Walker et al., 2018). Whereas modeling of tumor-immune dynamics has been a popular topic for some time (see the reviews in (Eftimie et al., 2011, 2016)), the number of such models that include *pro*-tumor immune effects are limited. den Breems and Eftimie (2016) incorporated M1 and M2 type macrophages in a 6-dimensional ODE model of tumor immune dynamics which included phenotypic switching between *anti*-tumor M1 macrophages and *pro*-tumor M2 macrophages. A more refined model of macrophage phenotypic plasticity was included in a more recent paper (Eftimie and Eftimie, 2018) concerning tumor-immune dynamics in the presence of an oncolytic virotherapy. Wilkie and Hahnfeldt (2017) have also developed a model of tumor-immune interactions that includes *pro*-tumor immune effects by including an immune-dependent carrying capacity for the tumor population. While a few models include the protumor effect of the immune response, this effect has not yet been included in a mathematical model for metastasis, as we do here.

### 1.3. Paper Outline

Section 2 is devoted to the development and basic analysis of the two-site model of tumor-educated immune mediated metastasis, including a subsection on parameter estimation and confrontation of the model to experimental data (Section 2.3). Once the model has been introduced and parameterized, we use it in Section 3 to perform three numerical experiments: simulations of primary resection, immunotherapy, and injury at a secondary site are shown. Model simulations demonstrate that tumor “education” of immune cells can significantly impair the effectiveness of immunotherapies and provide a potential explanation for rapid metastatic growth at the sites of injuries. We conclude with a discussion of our results and conclusions in Section 4.

## 2. Two Site Model of Immune-Mediated Metastasis

In this section, we describe our model for tumor-immune interactions at two anatomically distant sites. The modeling assumptions and the model itself are described in Section 2.1. We present the steady states of the model, including results concerning stability, in Section 2.2, and Section 2.3 introduces the functional coefficients and the parameter values used in the simulations that are the focus of Section 3. The section concludes with a comparison of our parameterized model predictions with experimental data from Kaplan et al. (2005).

### 2.1. The Model

Let us assume that there are two tumor sites of interest: the *primary* site, where the initial tumor develops, and a *secondary* site where a metastatic tumor will establish and grow. At both the primary and secondary sites (subscripts *i* = 1, 2, respectively) we model the time dynamics of four local cell populations: tumor cells, *u*_*i*_(*t*), necrotic cells, *v*_*i*_(*t*), cytotoxic (CT) immune cells, *x*_*i*_(*t*), and tumor-educated (TE) immune cells, *y*_*i*_(*t*). As we are modeling metastatic spread, the tumor cells of interest are those that are highly tumorigenic. Consequently, the tumor cells in our model can be interpreted as cancer stem cells (CSCs) or cells possessing the metastatic phenotype (see Section 1). In any case, we assume that a fraction, 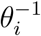, of the tumor cells are capable of metastasizing. The full model is depicted graphically in Figure 2 and in Equations (1)–(8). The time-evolution of the eight quantities of interest in our model is governed by the following system of equations:

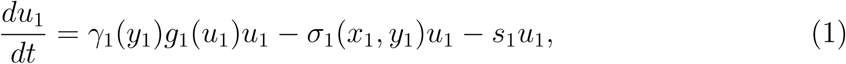

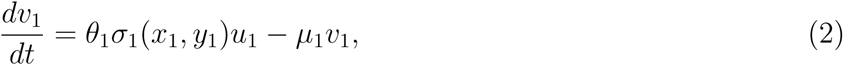

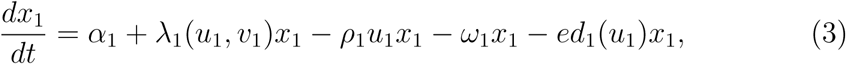

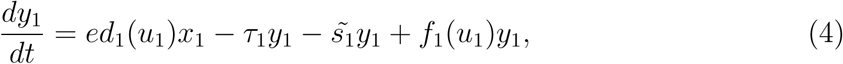

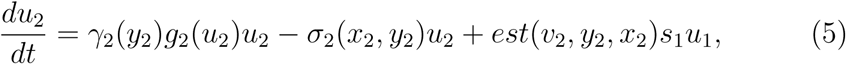

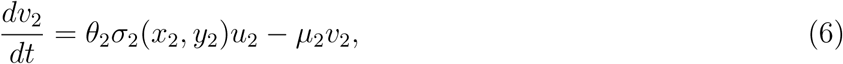

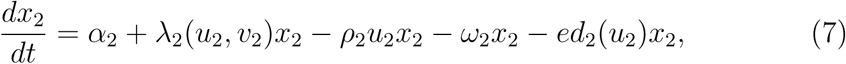

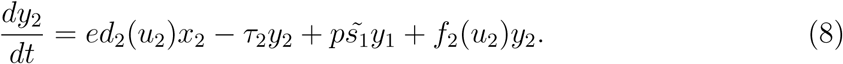

**Figure 2:**
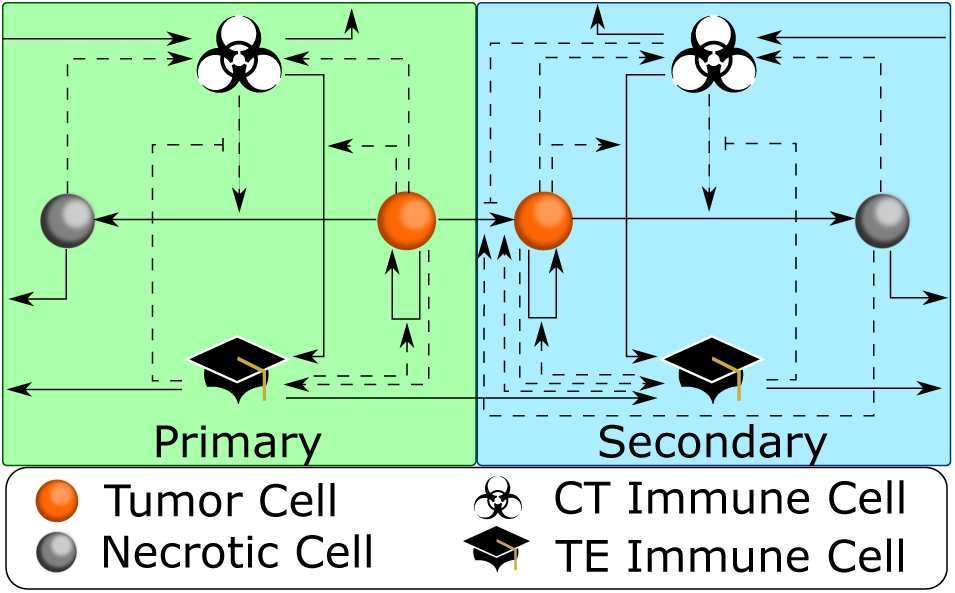
Cartoon model of the 8 ODE model of metastasis — Equations (1) – (8). Arrows indicate *positive* effects, and flat ends indicate *inhibitory* effects. Solid lines represent *direct* effects and dashed lines denote *indirect* influence. See text for details. Color figure available online.

The equations above incorporate the following biological assumptions (for details and references, please see Sections 1 and 2.3):

- In the absence of any immune cells, both tumor cell populations, *u*_*i*_ (*i* = 1, 2), proliferate according to the density-dependent growth rates, *g*_*i*_(*u*_*i*_), and perish at some non-negative rate *σ*_*i*_(*x*_*i*_, *y*_*i*_), thereby giving rise to necrotic cells, *v*_*i*_. CT immune cells, *x*_*i*_, can increase this tumor cell death rate, while TE immune cells, *y*_*i*_, can inhibit this CT immune response. Hence *σ*_*i*_(*x*_*i*_, *y*_*i*_) is *decreasing* in TE immune population, *y*_*i*_, and *increasing* in CT immune population, *x*_*i*_. In addition to their ability to suppress CT immune activity, TE immune cells can also stimulate tumor growth according to the increasing, bounded functions γ_*i*_(*y*_*i*_).
- Tumorigenic tumor cells are shed from the primary tumor into the surrounding vasculature proportionally to the primary tumor size with rate *s*_1_. A fraction, *est*(*v*_2_, *x*_2_, *y*_2_), of these cells will successfully navigate the blood stream, arrive at the secondary location, and contribute to the development of a metastatic tumor. The fraction of such cells depends on the local immune populations at the secondary site, *x*_2_ and *y*_2_, in addition to the necrotic cells populating the secondary site, *v*_2_. The fraction of successful cells, *est*(*v*_2_, *x*_2_, *y*_2_), is *increasing* in the TE immune cells, *y*_2_, and necrotic cell populations, *v*_2_, and *decreasing* in the local CT immune cell population, *x*_2_. We assume that establishment is more likely in the presence of necrotic cells (Shahriyari, 2016), but not impossible in their absence.
- At both sites (*i* = 1, 2) necrotic cells arise as a consequence of tumor cell death, and are lysed at rate *µ*_*i*_. Assuming that the *u*_*i*_ describe only a fraction of the total tumor burden, we include necrotic cells arising from the death of non-tumorigenic tumor cells by using the factors *θ*_*i*_.
- In addition to natural CT immune cell influx rates, *α*_1,2_, both local tumor cells and necrotic cells induce CT immune responses, described by the functions *λ*_*i*_(*v*_*i*_, *x*_*i*_), which are increasing in both arguments. CT immune cells perish at rates *ω*_*i*_, and are killed in interactions with tumor cells with rates *ρ*_*i*_. Finally, the local tumor population can induce a phenotypic transition of *anti*-tumor CT immune cells into *pro*-tumor TE immune cells. This “education” of immune cells is described by the increasing functions *ed*_*i*_(*u*_*i*_), *i* = 1, 2.
- In the absence of a tumor population at the primary site, there will be no TE immune cells. However, once a tumor is established at the primary site, TE immune cells can accumulate at the primary site in two ways: (1) by means of a tumor-induced phenotypic transition between CT and TE immune cell populations, and (2) by direct tumor recruitment of *pro*-tumor immune cells governed by the function *f*_*i*_(*u*_*i*_), *i* = 1, 2. The TE immune population at the primary site can decrease through natural death at rate *τ*_1_, or through loss into the circulatory system at rate 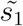
- A fraction, *p*, of those TE immune cells shed from the primary site arrive at the secondary site to supplement the previously described methods of TE immune cell accumulation — namely tumor “education” of CT immune cells and tumor-mediated recruitment. TE immune cells at the secondary site perish at rate *τ*_2_.
- We have assumed that the only significant shedding events occur from the primary site, a choice justified by previous theoretical work showing that shedding from the secondary site had negligible effects on the observed dynamics (Hartung et al., 2014).

### 2.2. Steady States

We quickly summarize the steady states of model (1)-(8) and their stability, without presenting the details of the analysis. Three different steady state expressions characterize the model:

1. A disease-free steady state, given by

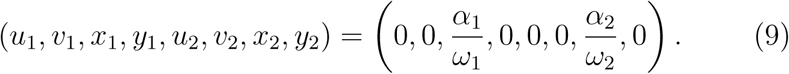
2. A metastatic-only steady state, given by

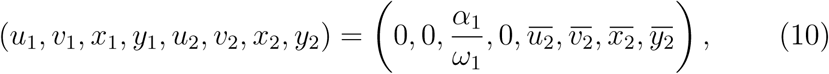

where the barred values (when they exist) are defined by the following equations:

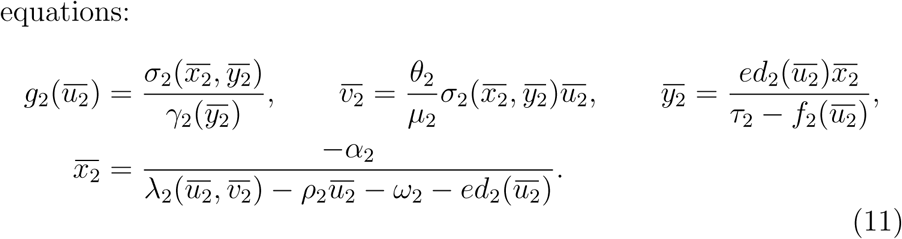
3. And a full-disease steady state expression, given by

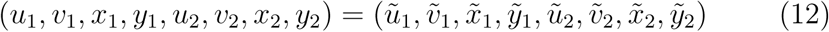

where the values on the RHS (when they exist) are defined by the following equations,

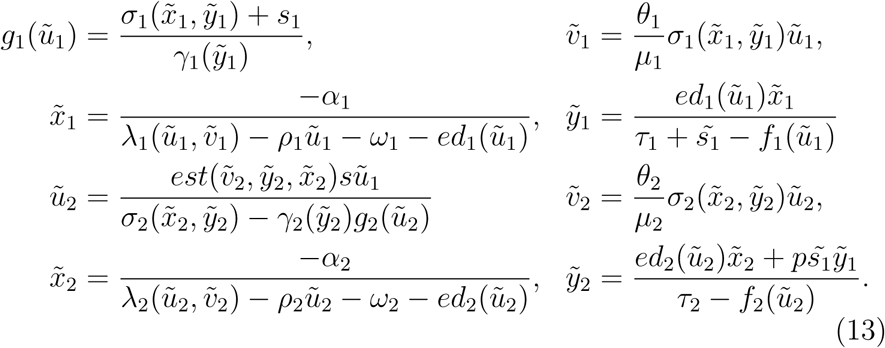

Representative solutions of the model illustrating the three different steady states are shown in Figure 3. Explicit conditions to ensure the stability of the disease-free and metastatic-only steady states are obtained, and presented in the following proposition.

**Figure 3:**
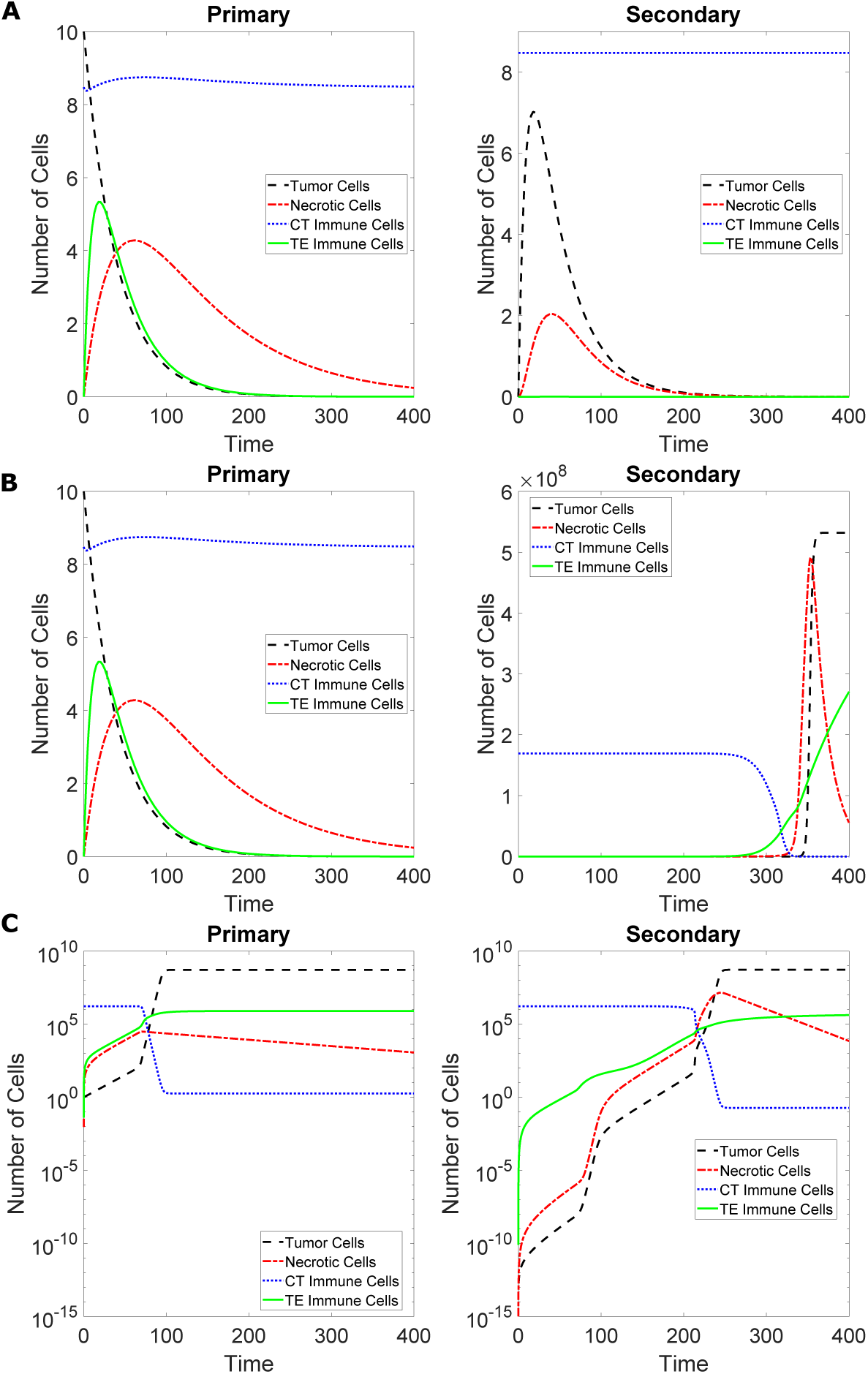
Solutions of the model (1) – (8) illustrating the three qualitatively different steady states. Left column: dynamics at the primary site. Right column: dynamics at the secondary site. Colors as indicated. Convergence to **A**: disease-free steady state. **B**: metastatic-only steady state. **C**: full disease steady state. Parameters from Table 1 used in **C**, and appropriately modified parameters used in **A** and **B** according to the conditions in Proposition 2.1. Color figure available online.

#### Proposition 2.1. *If*

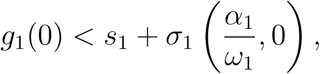

*then extinction of the primary tumor is stable.Further, the disease-free steady state is stable if and only if both*

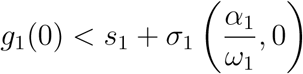

*and*

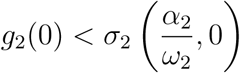

*are satisfied.*

#### Remark 2.2

*Note that the expressions for many of the non-trivial steady states have denominators which could potentially change signs. In order for these values to be biologically relevant, we insist on non-negativity of all the steady state expressions. In particular, the denominators cannot be allowed to change signs in our domains of interest. Using the literature-derived parameter estimates in Table 1 this requirement is satisfied.*

**Table 1:** Model Parameters and the values used in presented simulations.

| Parameter | Description | Value | Units | References |
| --- | --- | --- | --- | --- |
| $r_{1,2}$ | tumor growth rate | 0.38 | 1/day | Poleszczuk et al. (2016) |
| $K_{1,2}$ | tumor carrying capacity | $5.3196 \times 10^8$ | cells | Poleszczuk et al. (2016) |
| $\theta_{1,2}$ | CSC scaling constant | 65.67 | — | Enderling (2015);<br>Rhodes and Hillen (2016) |
| $\mu_{1,2}$ | dead cell lysis rate | 0.01, 0.05 | 1/day | Eikenberry et al. (2009);<br>Robertson-Tessi et al. (2015) |
| $\alpha_{1,2}$ | CT immune influx rate | $1 \times 10^6$ | 1/day | Eikenberry et al. (2009) |
| $\rho_{1,2}$ | fatal immune-tumor interaction rate | 0.001, 0.01 | 1/day | Eikenberry et al. (2009) |
| $\omega_{1,2}$ | CT decay rate | 0.59 | 1/day | Poleszczuk et al. (2016) |
| $\chi_{1,2}$ | immune education rate | $5 \times 10^{-5}$ | 1/day | den Breems and Eftimie (2016);<br>Kim et al. (2017) |
| $\tau_{1,2}$ | TE decay rate | 0.05 | 1/day | Eikenberry et al. (2009) |
| $s_1$ | tumor shedding rate | 0.01 | 1/day | Gupta and Massague (2006);<br>Joyce and Pollard (2009) |
| $\tilde{s}_1$ | TE shedding rate | 0.05 | 1/day | Eikenberry et al. (2009) |
| $p$ | proportion successful TE | $1 \times 10^{-4}$ | — | — |
| $max_{1,2}$ | max (increase) TE growth | 0.5 | — | — |
| $low_{1,2}$ | growth activation | 25424 | cells | — |
| $up_{1,2}$ | growth saturation | 110169 | cells | — |
| $lowD_{1,2}$ | death activation: TE | 25424 | cells | — |
| $upD_{1,2}$ | death saturation: TE | 110169 | cells | — |
| $minC_{1,2}$ | min death rate | 0.2 | 1/day | Orlando et al. (2013);<br>Saidel et al. (1976) |
| $maxC_{1,2}$ | max increase death | 0.1 | 1/day | — |
| $lowC_{1,2}$ | death activation: CT | 254237 | cells | — |
| $upC_{1,2}$ | death saturation: CT | 1101695 | cells | — |
| $a_{11,12}$ | CT expansion: tumor | 0.524 | 1/day | Kuznetsov and Knott (2001) |
| $a_{21,22}$ | CT expansion: dead | 0.786 | 1/day | — |
| $b_{11,12,21,22}$ | immune damping (dead;tumor) | $1.61 \times 10^5$ | cells | Kuznetsov and Knott (2001) |
| $a_{31,32}$ | TE expansion rate | 0.04 | 1/day | Kuznetsov and Knott (2001);<br>Poleszczuk et al. (2016) |
| $b_{31,32}$ | TE expansion damping | $1.6 \times 10^5$ | cells | Kuznetsov and Knott (2001);<br>Poleszczuk et al. (2016) |
| $max_{CT}$ | max (increase) establish rate | 100 | — | Gorelik (1983) |
| $lowEst_{CT,TE}$ | activation level: establish | 254237, 25424 | cells | — |
| $upEst_{CT,TE}$ | saturation level: establish | 1101695, 110169 | cells | — |
| $minEst_{TE}$ | min establish rate | 0.001 | 1/day | Cameron et al. (2000);<br>Chambers et al. (2002);<br>Joyce and Pollard (2009);<br>Mehlen and Puisieux (2006) |
| $maxEst_{TE}$ | max establish rate (increase) | 0.002 | 1/day | — |
| $low_V$ | activation: dead cells | $2.66 \times 10^7$ | cells | — |
| $up_V$ | saturation: dead cells | $2.93 \times 10^8$ | cells | — |
| $min_V$ | min establish rate | 0.001 | 1/day | — |
| $max_V$ | max establish rate (increase) | 0.999 | 1/day | — |

### 2.3 Parameter Estimation

Numerical exploration of the model necessitates that certain choices are made for the general functional coefficients in the model (1)–(8). In this section, we make our choices and parameterize the resulting model. As a simplifying generalization, we have assumed, as have others (Poleszczuk et al., 2016), that many of the model parameter values are shared between primary and secondary sites. This assumption is almost certainly incorrect (Hanin and Rose, 2018), but as a first approximation we contend it suffices. Table 1 summarizes the parameter values used in this paper together with appropriate references (where applicable).

- Tumor cell growth rates, *g*_*i*_(*u*_*i*_), *i* = 1, 2, are chosen to be of logistic type,

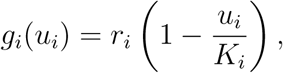

where *r*_*i*_ and *K*_*i*_ are growth rates and carrying capacities at sites *i* = 1, 2, respectively. Whereas we recognize that logistic growth is not the ideal choice for modeling tumor growth dynamics in the metastatic setting (Hartung et al., 2014), we have chosen to assume logistic growth to mirror the choices of other investigators (den Breems and Eftimie, 2016; Eftimie and Eftimie, 2018; Kuznetsov et al., 1994; Poleszczuk and Enderling, 2016). In the simulations presented below, we have used tumor growth rates and carrying capacities determined in Kuznetsov and Knott (2001) by fitting experimental data, and which were used again in more recent work in the setting of *abscopal* effects (Poleszczuk et al., 2016).
- Tumor cell death rates, *σ*_*i*_(*x*_*i*_, *y*_*i*_), are chosen to be a product of “switch-like” hyperbolic tangent functions as used by Olobatuyi et al. (2017). We include both an *increasing* version:

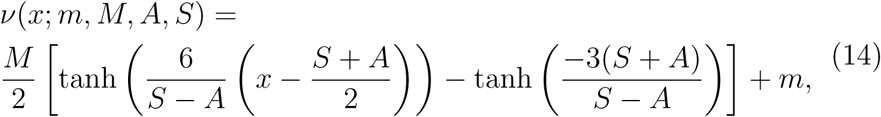

which increases from *m* at *x* = 0 to *m* + *M* as *x → ∞*, and a *decreasing* version:

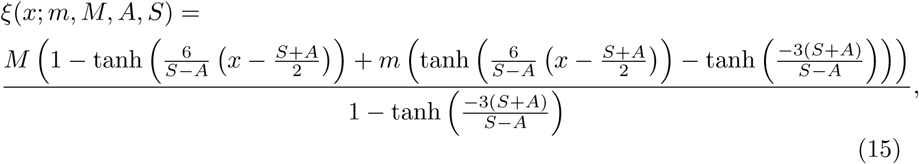

which decreases from *m* + *M* at *x* = 0 to *m* as *x → ∞*. Both of these “switch-like” functions can be specifically tuned using four parameters: activation (*A*) and saturation (*S*) thresholds together with lower (*m*) and upper (*m* + *M*) bounds on the domain [0, *∞*). The tumor cell death rates are then chosen as the product

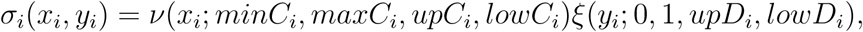

which increases in the CT immune cell populations, *x*_*i*_, and decreases in the TE immune cell populations, *y*_*i*_, *i* = 1, 2. The only parameter in this function that we were able to estimate from the literature is the minimum death rates, *minC*_*i*_, with the estimate coming from previous theoretical investigations (Orlando et al., 2013; Saidel et al., 1976). All remaining parameters were estimated conservatively, for example, most CT immune cell thresholds were chosen to be 15% (activation) and 65% (saturation) of the disease-free steady state value of CT immune cells 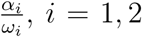, which was chosen to be of the order 10^6^ cells by tuning the parameters *α*_*i*_ (den Breems and Eftimie, 2016; Negus et al., 1997; Steidl et al., 2010)), and TE immune cell thresholds were then chosen to be an order of magnitude *lower* than those for CT immune cells.
- To model tumor and necrotic cell mediated immune cell expansion, we use the successful Michaelis-Menten type function which is a popular choice in tumor-immune models (Eftimie et al., 2011, 2016; Kuznetsov et al., 1994; Poleszczuk et al., 2016). As a consequence of their ubiquity, estimates have been made by several authors for the associated CT immune cell parameters. We have assumed that CT immune cell recruitment by tumor cells and necrotic cells is *additive*, i.e.

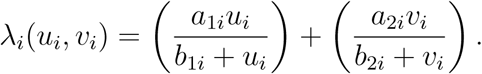

We have also included tumor-mediated recruitment of TE immune cells using the standard Michaelis-Menten type functions,

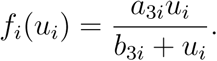

Whereas CT immune cell related parameters were easily found in the literature, no such estimates exist (to the authors’ knowledge) for TE immune cells. As a consequence, we have estimated the TE immune parameters by scaling the corresponding CT immune cell parameters by up to an order of magnitude.
- Due to its relatively recent experimental discovery, there has been little work done attempting to elucidate the precise mechanisms underlying the tumor-mediated phenotypic plasticity, or “education”, of CT immune cells. As a result, the only relevant literature from which we can inform our model is the theoretical work of den Breems and Eftimie (den Breems and Eftimie, 2016), in which the authors use mass-action kinetics to describe the tumor “education” of CT immune cells. Following this approach allows us to choose

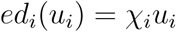

for some non-negative rate constants *χ*_*i*_, *i* = 1, 2. In the absence of additional evidence, we have chosen to use den Breem and Eftimie’s “polarization” rate as our “education” rate (den Breems and Eftimie, 2016; Kim et al., 2017). For further discussion, see Section 4.
- To model the TE immune cell enhancement of tumor growth, we have used the increasing hyperbolic tangent functions (14)

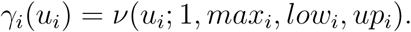

Thresholds were chosen as discussed previously, and we have assumed that TE immune cells could increase the tumor cell growth rate by at most 50%.
- Finally, we choose a model for the establishment of circulating tumor cells at the secondary site. Based on the evidence discussed in Section 1 we use

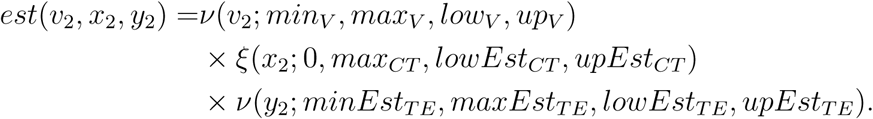

Immune cell thresholds were chosen as above, while the necrotic cell thresholds were chosen to be 5% and 55% of the tumor carrying capacities *K*_*i*_. Estimates of the rates involved have been informed by both previous experimental evidence (Cameron et al., 2000; Chambers et al., 2002; Gorelik, 1983; Joyce and Pollard, 2009; Mehlen and Puisieux, 2006) and the authors’ estimates.

The above discussion is summarized in Table 1, where the parameter values used in Section 3 are summarized with references (where applicable).

The initial conditions for all presented results (with the exceptions of Figure 3 A and B, where the number of tumor cells at the primary site as chosen to be larger for purposes of illustration) were chosen to be

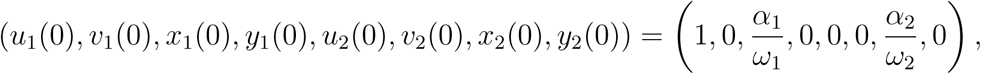

representing a slight perturbation of the disease-free steady state in which a single tumor cell has developed at the primary site. This choice allows for the inclusion of a time-dependent source of circulating tumor cells coming from the growing primary tumor — dynamics that are often neglected in injection (Cameron et al., 2000) or simulation (Eikenberry et al., 2009; Walker et al., 2018) studies.

As an initial validation of both the model and the chosen parameter values, we compare the calibrated model’s predicted dynamics at the secondary site with those observed experimentally by Kaplan et al. (2005). Following intradermal injection of 2 *×*10^6^ Lewis lung carcinoma (LLC) cells into murine flanks, Kaplan and colleagues measured the proportions of *pro*-tumor BMDCs and tumor cells at the metastatic site (lungs) at several time points (Figure 4, **A** (Figure 1 c in Kaplan et al. (2005))). As can be seen in Figure 4 (**A**) pro-tumor BMDCs arrived at the site of future metastasis well before the arrival of any tumor cells. Our model successfully captures this phenomenon, as seen in Figure 4 (**B**), with pro-tumor TE immune cells arriving at the future metastatic site in advance of any significant tumor colonization. Furthermore, our model accurately captures the approximate scales of this colonization, both in terms of *magnitude* (peaks of approximately 30% and 10% for BMDCs and tumor cells, respectively) and *timing* (approximately 2 weeks from initial colonization to tumor cell takeover).

**Figure 4:**
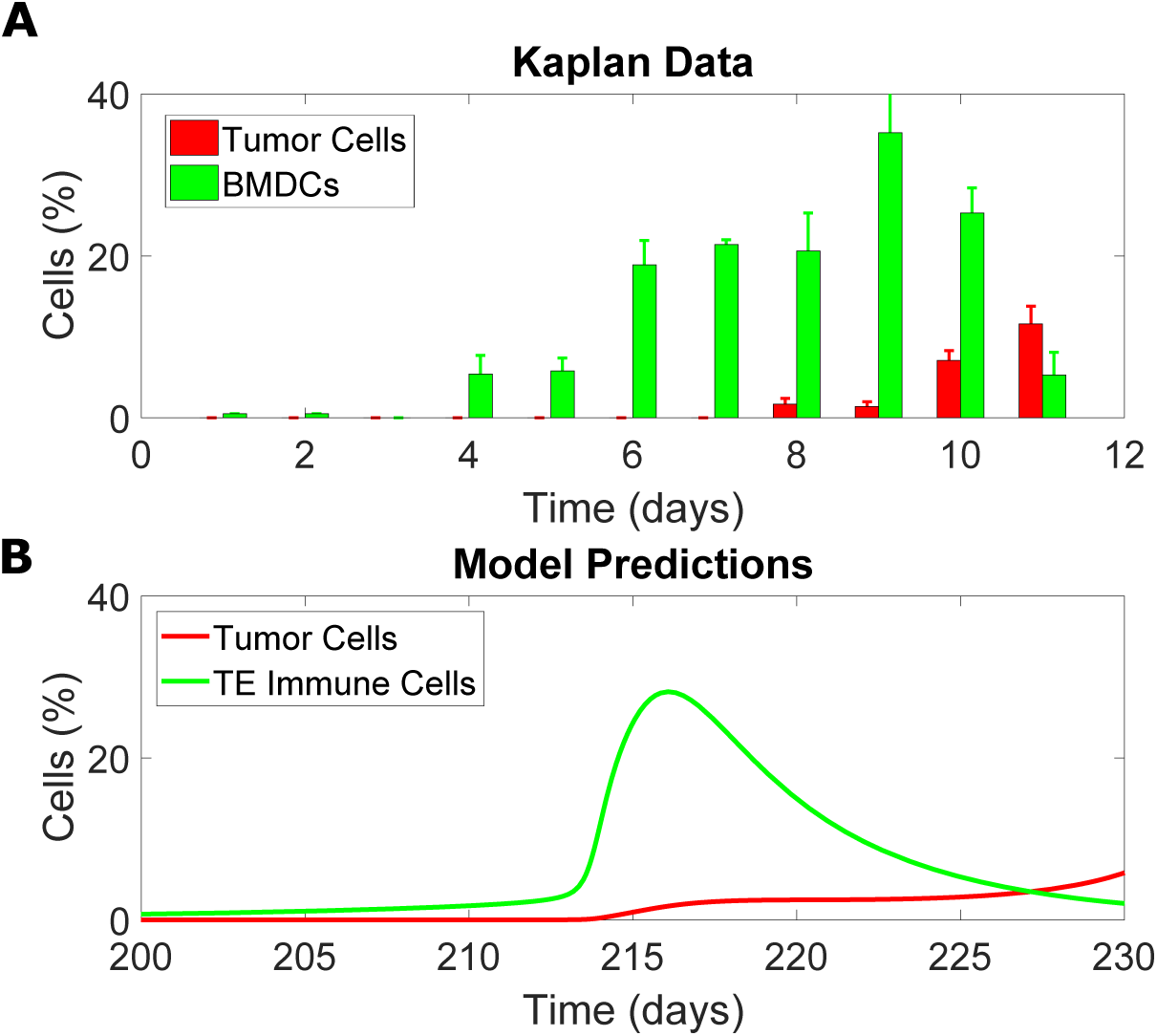
Comparison of experimental data from Kaplan et al. (2005) (**A**) to the model predicted dynamics at the secondary site (**B**). Time in the top plot is measured from the time of injection of 2 *×* 10^6^ LLC cancer cells, whereas time in the bottom plot is measured from the beginning of the simulation (primary tumor inception). In both cases, green corresponds to *pro*-tumor immune cells (BMDCs at top, and TE immune cells at bottom) and red corresponds to tumor cells. Color figure available online. **A** adapted from Kaplan et al. (2005), Figure 1 c.

There are, however, two shortcomings of this comparison. First, in our simulation, the primary tumor reaches a size of approximately 2 *×* 10^6^ cells (matching the size of the injection used by Kaplan et al. (2005)) after 84 days, meaning that the delay to the dynamics presented in Figure 4 is approximately 120 days. While this shortcoming may appear problematic, it may simply be a consequence of the differences between the details of the experiment and the simulation. Second, the *shape* of the pro-tumor immune curve in the experimental data — and the location of the peak, in particular — is not well approximated by our simulation results, with the simulated peak occurring much *earlier* than the experimentally observed peak. Although these are both important shortcomings, we note that the data from Kaplan et al. (2005) was not used in the calibration of our model. Consequently, some inconsistencies may be reasonably expected, and we contend that the successes described previously — those of *order* of arrival and general scales being well approximated — are sufficient to claim that the parameters in Table 1 are biologically feasible and that the qualitative results of the calibrated model reflect the true biology.

## 3. Model Simulations

Now that we have used experimental evidence and previous literature-derived estimates to specify the model parameters, and we have confirmed that these parameters can accurately reproduce experimental results (Figure 4), we perform three clinically relevant numerical simulations of the model in order to further investigate the implications of the immune-mediated theory of metastasis. This section investigates the effects of primary resection, immune therapies, and injury at the secondary site on disease progression.

### 3.1. Primary Resection

When possible, surgical removal (resection) of a tumor can be the preferred method of treatment. Unfortunately, this treatment is not always effective and may only offer temporary relief, with local recurrence or metastatic disease appearing after a short period of apparent health. Our model framework allows us to interrogate the effect of primary resection on the dynamics of the secondary tumor. We study two cases. In the **first case**, we assume that the disease free equilibrium is locally unstable. In this case, each resection of less than 100% efficiency leads to recurrence of the tumors. Here we are interested in the time delay before re-growth occurs. Figure 5 shows the tumor cell dynamics at the primary (left) and secondary (right) sites using the parameters from Table 1. The untreated dynamics at both sites are represented by the black curves. A primary tumor develops relatively quickly and reaches the local carrying capacity after approximately 100 days, with the secondary tumor only fully developing approximately 150 days later (notice the different time intervals shown on the horizontal-axis).

**Figure 5:**
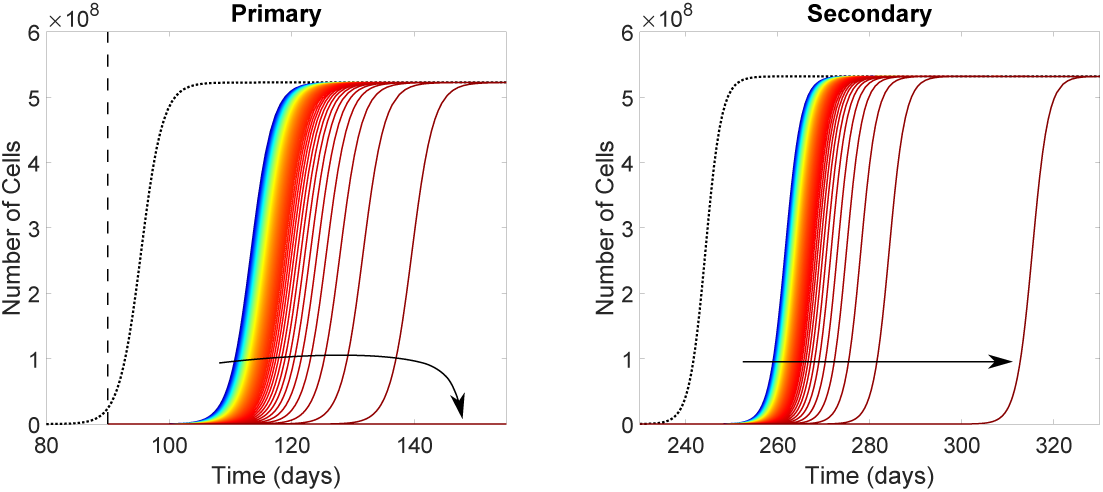
Effects of primary resection on tumor population dynamics at the primary (left) and secondary (right) sites. Primary tumor was removed at time *t* = 90 days (vertical dashed line in left plot) with efficiency ranging from 99.99% to 100% (blue to red). Arrow indicates the direction of increasing resection efficiency. The black dotted curves represent control dynamics. Parameters as in Table 1. Color figure available online.

Note that the saturation observed at both sites can be explained in terms of CSCs. Indeed, we have assumed that the quantities *u*_1_ and *u*_2_ represent *tumorigenic* cells within the tumor populations at the primary and secondary sites, respectively. Therefore, saturation may correspond to the homeostatically stable population of CSCs, and may not necessarily represent the end of tumor growth. As the fraction of CSCs within a tumor population is a hotly debated topic (Enderling, 2015), with a number of theoretical results suggesting that pure CSC tumors are possible (Rhodes and Hillen, 2016), this explanation is not unfounded. Alternatively, the rapid saturation at the primary site may suggest that, in this case, the subject would succumb to the primary tumor before the advent of any significant metastatic disease.

We have simulated primary resection by removing a specified fraction — the *resection efficiency* — of *all* populations at the primary site at time *t* = 90 days (vertical dashed line in left plot of Figure 5) (Eikenberry et al., 2009). The resection time was chosen such that the primary tumor grew sufficiently large so that it could be detected by clinicians (order 10^7^ cells (Friberg and Mattson, 1997; Eftimie and Eftimie, 2018)). Resection efficiencies in Figure 5 range from 99.99% (blue) to 100% (red). As expected, increasing the resection efficiency increases the delay in both local recurrence and metastasis. Using the parameters in Table 1, no resection efficiencies *below* 100% can prevent local recurrence, and no resection efficiencies can prevent metastasis.

In a **second case** we assume that tumor extinction at the secondary site is stable by reducing the tumor growth rate *r*_2_ (see Proposition 2.1). Then the model exhibits more realistic bistable behavior at the expense of much slower metastatic growth. The secondary tumor dynamics in response to 100% efficient primary resection are presented in Figure 6 for varying times of primary resection. As a control, we present the secondary tumor dynamics in the absence of any primary intervention as the black curve. A consequence of primary resection is that the secondary site loses a source of tumor cells. If this primary intervention occurs sufficiently early, the secondary tumor is too small to support itself, resulting in metastatic extinction (green curves in Figure 6).

**Figure 6:**
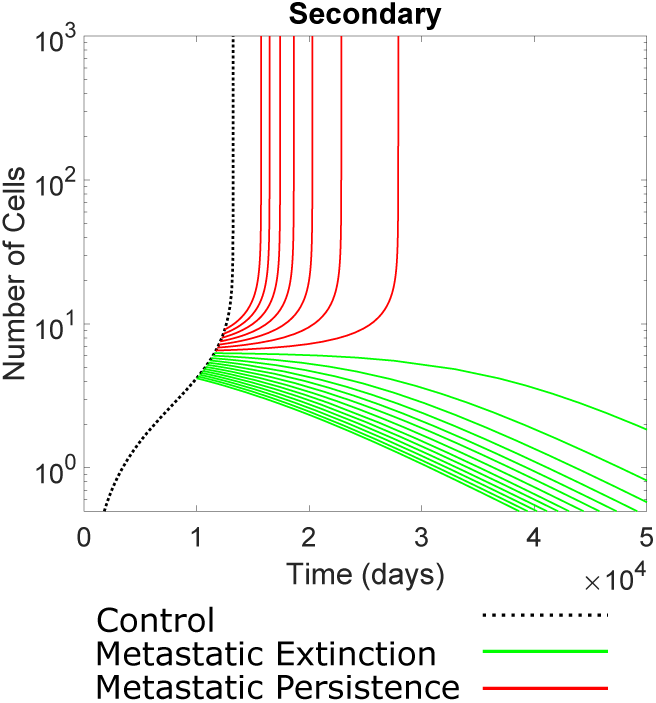
Effect of primary resection on secondary tumor dynamics for various times of primary resection, ranging between *t* = 10000 and *t* = 12500 days. Primary resection was 100% efficient, meaning there was no influence on secondary site from the primary site following resection. Secondary tumor growth rate was *r*_*2*_ = 0.2999 so that extinction of metastases was stable. The black trajectory shows secondary tumor growth without primary resection and acts as a control curve. Green trajectories are destined for extinction, while red trajectories are destined for full secondary tumor. Color figure available online.

On the other hand, if enough time has passed with the primary tumor present to ensure that the metastatic tumor is large enough so that it can maintain growth even in the absence of the source of cells from the primary tumor, then we observe rapid metastatic growth, possibly following a period of dormancy (red curves in Figure 6). Two important observations should be made in this case. First, the final metastatic tumor density is smaller when compared to the control case, and second, primary resection can trigger an extended period of dormancy at the secondary site.

### 3.2. Immune Therapy

While there is a significant diversity of immunotherapeutic techniques in cancer treatment, they all share the same goal: increase the number or effectiveness of CT immune cells in order to elicit a strong antitumor response. The promise of harnessing the power of the immune system to treat tumors has inspired significant experimental and theoretical investigation. Unfortunately, in many cases, the promises of immunotherapy have not come to fruition, with relatively low response rates for both single (10% 30%) and combination therapies (50% 60%) (Emens et al., 2017). A potential explanation for this shortcoming may be found in the contradictory roles of the immune system in cancer progression (Section 1), but this possibility has remained largely unexplored. In this section, we consider the implications of immune phenotypic plasticity on the effectiveness of immunotherapies.

In the following, “immunotherapy” will be simplified to any intervention that results in an increased influx of CT immune cells. Therefore, increasing the CT immune cell influx rates, *α*_1_ and *α*_2_, by some scaling factor will serve as a simplified model of immunotherapy. As was the case for primary resection, therapeutic interventions can only be undertaken in the case that a primary tumor has been identified clinically, so we begin therapy at time *t* = 90 days and maintain the therapy until the end of the simulation. Under these conditions, the model predicts little effect on the dynamics at the primary site — not an unexpected result based on the low response rates of many immunotherapies — so we present only the dynamics at the secondary site; thereby shedding light onto the effects of immunotherapy on small, clinically undetectable metastatic tumors, which is particularly relevant to the clinical setting.

The model predicted results of immotherapy are presented in Figure 7. Figure 7 (left) shows the dynamics of the secondary tumor cell population for various scaling factors of the CT immune cell influx rates *α*_1_ and *α*_2_, with the scaling factor increasing from blue to red. Figure 7 (right) shows the time to half the carrying capacity (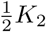 — horizontal dashed line in left plot) as a function of the scaling factor; in other words, the right plot shows the intersection times of the solution curves and the dashed line in the left plot. Of note is the non-monotonicity of the rightmost plot. For small increases to the immune influx rates we see significant improvement in tumor delay. However, there is an optimal increase factor, above which the effects of the immunotherapy are actually *detrimental* relative to the optimal and, if increased by a sufficiently large factor, we can have detrimental effects even compared to the control case (results not shown).

**Figure 7:**
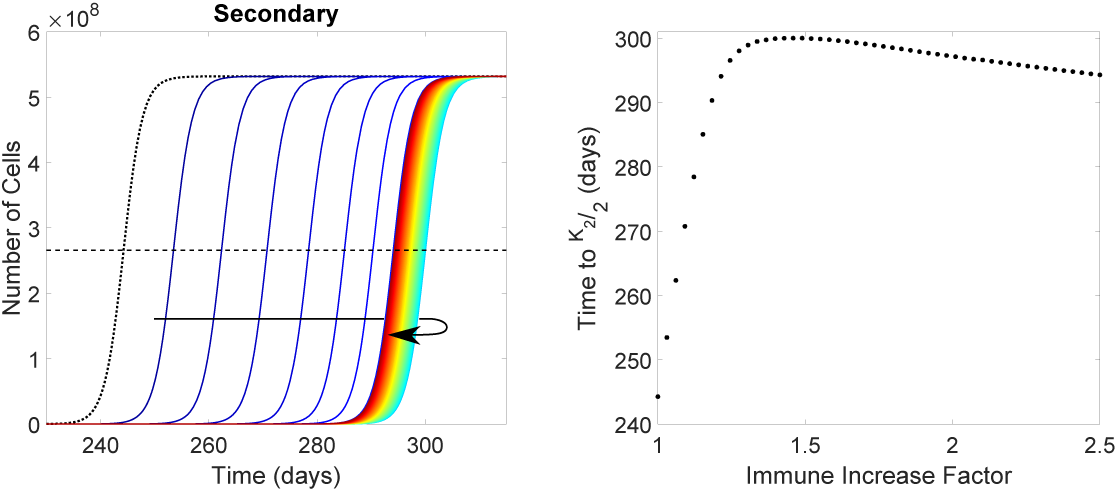
Effect of immune therapy on growth of the metastatic tumor. Left: Tumor cell dynamics at the secondary site for increasing CT immune cell influx rates, *α*_*1*_ and *α*_*2*_. Therapy administered beginning from time *t* = 90 days, and maintained over the course of the simulation. Values are increasing from blue to red (in direction of arrow). Leftmost (black dotted) curve represents control dynamics. Dash line represents half the carrying capacity,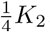. Right: Time secondary tumor reaches half its carrying capacity as a function of the factor the CT immune cell influx rate increased. Color figure available online.

In order to determine the mechanism responsible for the non-monotone dynamics in Figure 7, we simulate a modified version of the previous immunotherapy. In addition to the increased CT immune cell influx rate, we assume that our simulated immunotherapy is capable of *preventing* tumor education of CT immune cells (i.e. the education rates vanish: *χ*_1,2_ = 0). The model predicted effects of such an intervention are presented in Figure 8. As in the previous figure, the left plot shows the tumor cell dynamics at the secondary site for varying strengths of immunotherapy (increasing from blue to red) and the right plot shows the times our solutions reach the endpoint as a function of immunotherapy strength. Note that by preventing tumor education of CT immune cells, the resulting steady state tumor density at the secondary site is significantly diminished, so we use 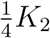 as our endpoint instead of the previously used 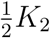

**Figure 8:**
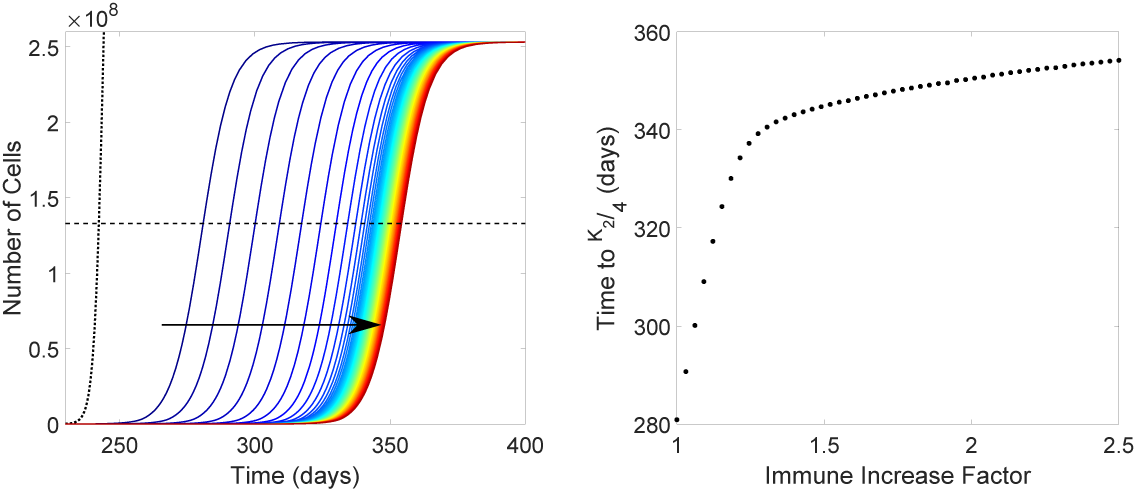
Effect of two-pronged immune therapy on growth of the metastatic tumor. Left: Tumor cell dynamics at the secondary site for increasing CT immune cell influx rates, *α*_*1*_ and *α*_*2*_, and prevention of immune education, *χ*_*1,2*_ = 0. Therapy administered beginning from time *t* = 90 days, and maintained over the course of the simulation. Values are increasing from blue to red (in direction of arrow). Leftmost (black dotted) curve represents control dynamics. Dash line represents one quarter the carrying capacity, 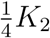 Right: Time secondary tumor reaches a quarter its carrying capacity as a function of the factor the CT immune cell influx rate increased. Color figure available online.

In contrast to the previous case, the rightmost plot in Figure 8 is *monotonically* increasing. Although there is a significant slowing of growth in the right plot, the time to endpoint continues to increase for all CT influx rate scalings tested. Noting, in addition, that the ability of the secondary tumor to directly recruit *pro*-tumor immune cells to the secondary site was not affected by our simulated therapy, it follows that the tumor-induced phenotypic plasticity between CT and TE immune cells is key in the non-monotonic dynamics of Figure 7.

### 3.3. Metastasis to Sites of Injury

Our modeling framework provides us the opportunity to investigate whether or not this theory of immune-mediated metastasis is sufficient to explain the observations of metastatic spread to sites of injury. We simulate an injury at the secondary site at time *t* by pausing the simulation at time *t*, adding 10^7^ cells to the necrotic compartment, and restarting the simulation with this adjusted initial condition. Evaluation of this injury’s effect on the secondary tumor dynamics is done by reporting the time when the secondary tumor reaches a population of 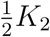 cells (referred to hereafter as the “endpoint”).

Figure 9 shows the time to endpoint as a function of the time that an injury at the secondary site is incurred. Control results are presented as the horizontal dashed line, so that times *above* this line are *beneficial* to patient survival (green), and those *below* the line are *detrimental* (red). The model predicts a clear distinction between *early* and *late* injuries. Injuries incurred earlier in the progression of the metastatic tumor are actually beneficial to the patient — delaying metastatic growth by up to nearly 4 months — whereas those that occur in the later stages of disease progression are detrimental to the patient, reducing the time to endpoint by up to two months.

**Figure 9:**
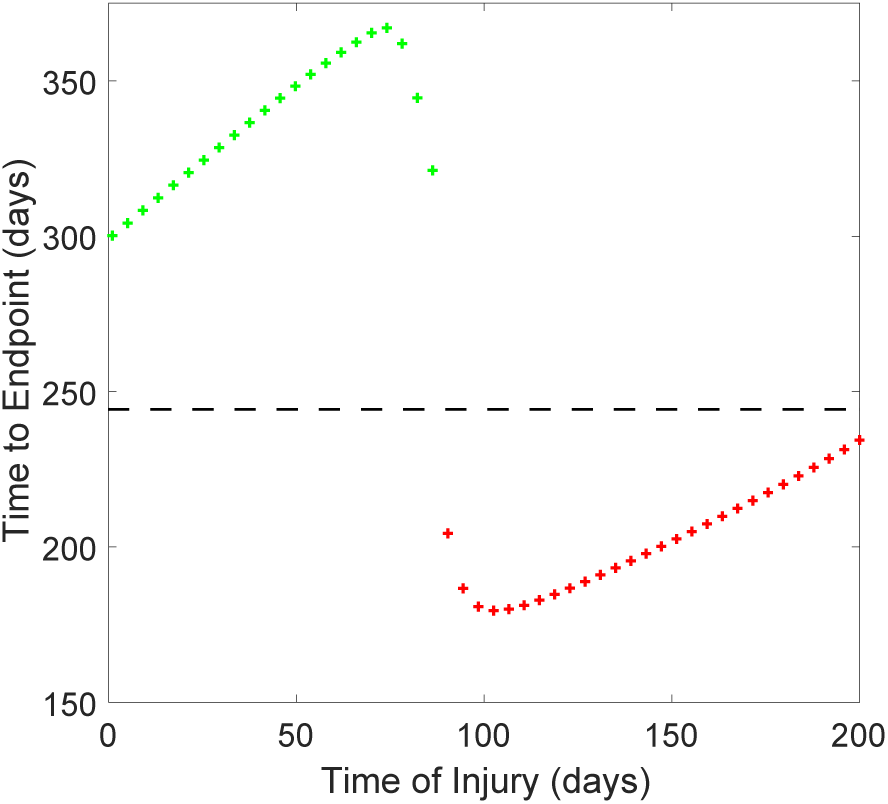
Time required for secondary tumor to reach 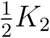 cells as a function of the time an injury of 1*×*10^7^ necrotic cells was incurred. Dashed line represents the control value. Green (above dashed line) indicates a desirable outcome, while red (below dashed line) indicates an undesirable outcome. Parameters as in Table 1. Color figure available online.

A glimpse into the mechanisms underlying the biphasic dynamics of Figure 9 are presented in Figure 10, where the dynamics of all cell populations at the secondary site are presented for three situations: control dynamics (black dotted curves), an early injury (green curves) and a late injury (red curves). Early and late injury times were chosen to be the times corresponding to the maximum and minimum times to endpoint from Figure 9, respectively (indicated by the colored arrows in Figure 10). Note that the early injury occurs slightly *before* clinical detectability of the primary tumor (which we have taken to be 90 days), while the late injury occurs slightly *after*.

**Figure 10:**
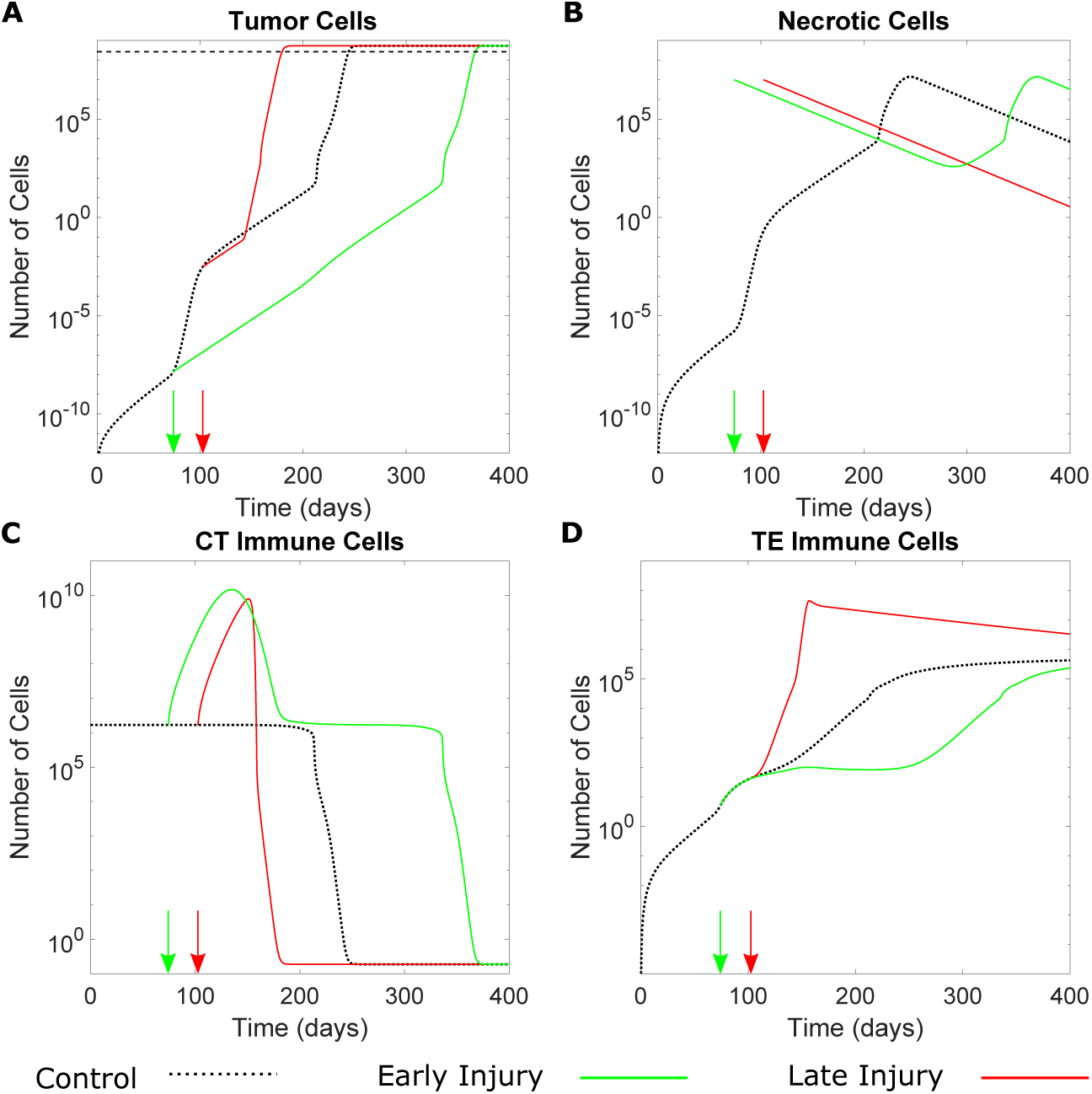
Dynamics of the tumor cells (**A**), necrotic cells (**B**), CT immune cells (**C**), and TE immune cells (**D**) at the secondary tumor site upon the simulation of an injury. Two injury times are presented (arrows): an *early* injury at *t* = 74.1 days (green) and a *late* injury at *t* = 102.5 days (red). Injury was 1 *×* 10^7^ necrotic cells. Dashed line in (A) represents endpoint value of 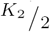 Dotted black curves in each plot denote control dynamics. Parameters as in Table 1. Color figure available online.

The dynamics in response to the *early* injury closely follow the traditional view of both injury response and tumor-immune dynamics: there is a robust, transient CT immune response to the injury (panel (C), green), which inhibits the phase of rapid tumor growth observed in the control dynamics beginning at approximately *t* = 75 days (panel (A), black). As a consequence of this slowed tumor growth, the TE immune population suffers an extended period of stagnation (panel (D), green), thereby slowing subsequent tumor growth and resulting in a significant delay in tumor progression at the secondary site.

In contrast, the dynamics in response to the *late* injury are remarkably different, and instead of delayed tumor growth, metastatic “blow-up” — rapid metastatic growth in response to en external stimulus — is predicted. Although the late injury induces a similar CT immune response, it is significantly foreshortened in comparison to the *early* injury case (panel (C)). Moreover, the TE immune population undergoes a period of rapid growth coinciding with the CT immune response (panel (D), red). Taken together, and noting that the tumor population has undergone a period of significant growth between the two injury times (panel (A), control), we conclude that the larger secondary tumor present at the time of the late injury more effectively corrupts, or educates, the CT immune response to the local injury. In fact, in simulations where the education rate was *decreased*, the injury no longer elicited a *pro*-tumor response (results not shown), demonstrating that tumor education of the CT immune response is vital to the dynamics reported in Figures 9 and 10.

The result of this education is a robust population of pro-tumor TE immune cells which stimulates rapid tumor growth much earlier than in the control case — that is to say that metastatic “blow-up” is observed. Therefore, our model predicts that rapid metastatic growth at the site of injury necessitates the presence of a sufficiently large local tumor population in order to adequately corrupt/educate the injury-induced CT immune response; otherwise, the immune response to injury is anti-tumor, as traditionally expected. Moreover, our results refine those of Eikenberry et al. (2009), whose mathematical model of metastatic melanoma suggested “blow-up” was the result of a depleted CT immune population. Indeed, “blow-up” in our model was a result of a decrease in the CT immune population *and* a corresponding increase in the TE immune population as a consequence of tumor “education.”

## 4. Discussion

There is a growing body of evidence implicating pro-tumor effects of the immune system in cancer development and metastatic spread (see the extended list of references in Section 1). Inspired by Shahriyari’s synthesis of this evidence (Shahriyari, 2016) to formulate a theory of metastasis in which the immune system plays a major role — which we have called the “immune-mediated” theory of metastasis (Cohen et al., 2015) — we have developed a mathematical model of tumor-immune interactions at two anatomically distant sites to interrogate the validity and the implications of this hypothesis.

Validation of our modeling approach and our literature-derived parameter estimates was done by confronting the model to experimental data from Kaplan et al. (2005). We found that the model correctly predicted the preparation of the PMN by protumor TE immune cells *prior* to the arrival of any tumor cells in addition to accurately reproducing the relative *magnitude* and *timing* of this PMN preparation (see Figure 4). It is important to note that these results were in the absence of any explicit fitting to the Kaplan data, and that the Kaplan data was not used to calibrate the model. Consequently, the discrepancies between data and model predictions are not be particularly concerning.

Once validated, the model was used to numerically explore the implications of the immune-mediated theory of metastasis. We simulated primary resection surgeries, immunotherapeutic interventions, and injuries at the secondary site. Metastasis is relatively robust in the face of primary resection, with metastatic extinction only possible in certain parameter regimes, and only if the primary tumor is completely removed sufficiently early (Figure 6). In response to the loss of cells arriving from the primary tumor, metastatic dormancy could be observed. A second set of numerical experiments concerned the effects of tumor-education on the efficacy of immunotherapies. We found that tumor-induced phenotypic plasticity between anti-and protumor immune cells provides a potential explanation for the relatively poor performance of many immunotherapies (Emens et al., 2017). Moreover, our model predicts that the most successful approach to improving the efficacy of immunotherapies is to inhibit tumor-induced phenotypic plasticity thereby allowing the CT immune cells to play their anti-tumor roles. This result has been recently demonstrated experimentally by Park et al. (2018), lending further credibility to the results presented herein.

Finally, we asked whether or not the immune-mediated theory of metastasis could provide an explanation for metastasis to sites of injury by simulating an injury at the secondary site. We found that the CT immune response elicited by an injury was *anti*-tumor in the absence of a significant metastatic tumor cell population at the secondary site, and *pro*-tumor if this population was sufficiently large to corrupt the incoming CT immune cells, forcing them to play a protumor role (Figure 10). Not only do these findings support the suggestion of Kumar and Manjunatha (2013) that a population of tumor cells is required at the injury site *prior* to the injury to see metastasis establish at that site, but they also suggest that tumor-induced phenotypic plasticity plays a crucial role in such establishment.

In the work above, we considered a secondary *site*, but the secondary tumor dynamics could also be interpreted as the total *metastatic burden* by appropriate choice of growth functions. Furthermore, we considered only one secondary site, but this could easily be extended to include *N* sites with anatomically motivated connection network as in (Poleszczuk et al., 2016; Franßen et al., 2018). We provide a brief sketch of such a model now. Let *u*_*i*_, *v*_*i*_, *x*_*i*_, and *y*_*i*_ denote the number of cancer, necrotic, CT, and TE cells at tumor site *i*, where *i* = 1, 2, *… N*. Let *ϕ*_*i,j*_, *ψ*_*i,j*_, and *ζ*_*i,j*_ denote the number of tumor cells, TE immune cells, and CT immune cells respectively, leaving site *j* and arriving at site *i*. We assume that necrotic cells do not travel between sites. Under the above assumptions, we arrive at the following *N* site model for tumor-immune interactions including both pro-and anti-tumor immune effects:

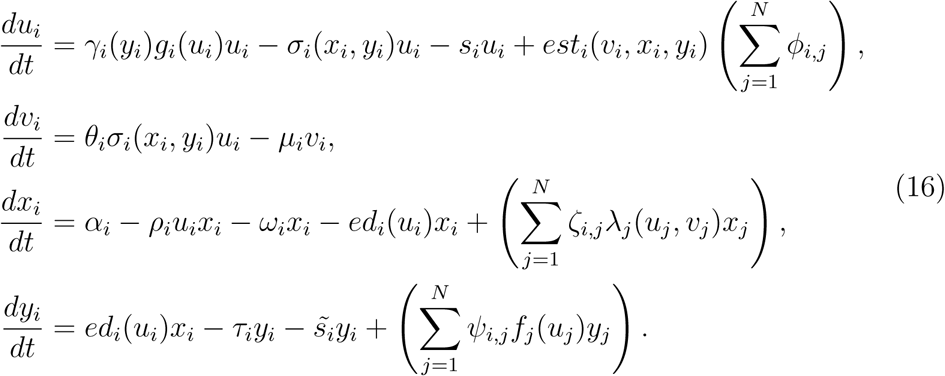

Modeling of the connection terms, *ϕ*_*i,j*_, *ψ*_*i,j*_, and *ζ*_*i,j*_, is a complicated problem (see the works of Poleszczuk et al. (2016); Walker et al. (2018, 2017) for discussion on the *ζ*_*i,j*_ term) and is left as future research.

Instead of complicating the present model further, it may also be of interest to simplify it in order to perform more rigorous mathematical analysis with the aim of discovering the mechanisms underlying the dynamics described in Section 3. Such simplification is the focus of a current study, with results forthcoming.

As the explicit inclusion of protumor immune cells in mathematical models for tumor-immune dynamics is relatively new, the model for phenotypic plasticity between immune types was chosen to follow simple mass-action kinetics which are most likely too simplistic. den Breems and Eftimie (2016), in their model of M1/M2 macrophages, also modeled the transition using mass-action kinetics. However, Eftimie and Eftimie (2018) have recently proposed a more sophisticated transition function. Based on the important effect that these phenotypic transitions appear to have on the overall dynamics (above, and in (den Breems and Eftimie, 2016; Eftimie and Eftimie, 2018)), research looking to uncover the underlying dynamics of this phenotypic plasticity is warranted.

The model developed here includes a significant number of parameters, with many of them not previously estimated. Consequently, the results we have presented above should be taken with some degree of caution. We do note that some of the TE immune related parameters — recruitment rate by the tumor, for example — may be underestimated (Oleinika et al., 2013), meaning that the observed effects may be conservative. Further experimental and theoretical work must be done in order to validate the predictions made herein before specific therapeutic recommendations can be made. Specializing the model to focus on specific immune cells in a particular cancer may provide more clinically relevant results, and is the focus of a current study.

Overall, our modeling approach showcases the importance of including protumor effects — and tumor-induced phenotypic plasticity in particular — in models of tumor-immune interactions. By confronting our mathematical model to experimental data, we performed meaningful simulations of complex biological phenomena, which provided important insight into the underlying dynamics; insight that may be obscured in traditional biological experimentation. We believe that our research can help inform the design of future experiments and clinical investigations focused on elucidating the precise nature of the protumor role of the immune system in cancer progression and metastasis.

## Acknowledgements

AR acknowledges support through an NSERC CGS-M scholarship and a PIMS Accelerator Award. TH acknowledges support through an NSERC Discovery Grant.

